# Nutribloods: novel synthetic lepidopteran haemolymphs for understanding insect-microbe interactions *in vitro*

**DOI:** 10.1101/2024.04.17.589926

**Authors:** Robert Holdbrook, Joanna L. Randall, Catherine E. Reavey, Awawing A. Andongma, Annabel Rice, Judith A. Smith, Stephen J. Simpson, Sheena C. Cotter, Kenneth Wilson

## Abstract

Understanding the role of nutrients in microbial population dynamics relies on a sound appreciation of their nutritional environment and how this may vary in different habitats. For microbial pathogens and commensals, this can be especially challenging because the microbe may share nutritional resources with its host. Here we design a series of 20 synthetic haemolymphs (*nutribloods*) that mimic haemolymph nutrient profiles of caterpillars fed on one of 20 chemically-defined diets, that vary in their protein:carbohydrate (P:C) ratio and caloric density. Using these, we are able to simulate the range of nutritional conditions that insect blood pathogens might face, providing a model system for understanding the role of nutrition in microbial growth. We tested this using the entomopathogen, *Xenorhabdus nematophila*, a gram-positive extracellular bacterium of insect hosts. This revealed that whilst bacterial fitness peaked in *nutriblood* nutrient space that was high in carbohydrates and low in proteins, levels of amino acids in the nutribloods also appear to be an important driving force for bacterial growth. Using synthetic haemolymphs that had average levels of all nutrients other than carbohydrate, protein or amino acids, we also established that bacterial growth is generally enhanced by carbohydrate and amino acids but reduced by proteins. Here, we have established a tractable model system for examining the role that nutrition plays in the growth of an entomopathogenic bacterium. In future work, this model host-pathogen system can be used to test a range of nutritionally-driven processes, including competition during co-infection and interactions with the host microbiome, as well as comparative studies of other entomopathogens.

## INTRODUCTION

All organisms require nutritional resources to thrive and they must adapt phenotypically and/or genetically to cope with variation in their nutritional environments (Kussell and Leibler 2005). For free-living organisms, this can often be relatively straightforward to quantify, but for organisms in symbiotic relationships this can prove more challenging because they may share nutritional resources with their symbiotic partner (Douglas 2021). Understanding these nutritional requirements may best be explored by studying the symbionts in isolation, where this is possible. Many microbes exist as parasitic, commensal or mutualistic symbionts of animal hosts and understanding their microbial nutritional requirements can aid our ability to exploit this to reduce the impact of damaging parasites or to facilitate beneficial commensal/mutualistic microbes. Indeed, there is strong evidence that animals may ‘self-medicate’ in response to parasitic infection by changing their diet in order to either improve their immune response or to limit the damage the parasite may cause (Erler et al. 2024; Huffman 2003; Singer, Mace, and Bernays 2009). Much of the current evidence for self-medication is anecdotal rather than experimental (Schmid-Hempel 2021). For example, it is assumed that the ingestion of tannin-rich vegetation by sifaka lemurs (Carrai et al. 2003), consumption of medicinal plants by Japanese macaques (Tasdemir et al. 2020) and porcupines (Viviano et al. 2022) are forms of self-mediation, but this remains to be tested experimentally. Experimental studies demonstrate that *Spodoptera* caterpillars increase the levels of protein in their diet to combat bacterial or viral infections (Povey et al. 2009; Lee et al. 2006; Povey et al. 2014), though identifying the precise mechanisms underpinning self-medication can prove difficult.

Nutritional Geometry (NG) is a state-space nutritional modelling approach geared towards the analysis of organismal nutritional requirements. A key aspect of this approach is the identification of an ‘intake target’ representing a nutrient balance that maximises fitness (Simpson and Raubenheimer 2012). So far, NG has been applied mainly to vertebrates and invertebrates of various taxa, revealing the role of the macronutrients, protein, lipids and carbohydrates in the optimisation of various fitness traits such as survival, fecundity and immunity (reviewed by Simpson & Raubenheimer, 2012). Microbes, such as bacteria, also become adapted to particular resource environments and may show poor performance on alternative resources (Litchman, Edwards, and Klausmeier 2015; Pulkkinen et al. 2018), hence microbes can also express intake targets (Dussutour et al. 2010). Although identification of nutritional optima is important for *in vitro* mass culture of microbes, studies investigating microbial nutrient-use usually involve batch cultures with generic media containing multiple nutrients that are varied simultaneously (e.g. (Pulkkinen et al., 2018)) or the modification of a single dietary component (Bowen et al. 2012; Kooliyottil et al. 2014).

The genera *Xenorhabdus* and *Photorhabdus* comprise gram-negative bacteria belonging to the family Enterobacteriaceae that have symbiotic relationships with entomopathogenic nematodes of the families *Steinernematidae* and *Heterorhabditidae*, respectively (Boemare, Akhurst, and Mourant 1993). These bacteria are widely used as models due to their dual role as mutualists and pathogens, their variable range of host species, and their importance for biological pest control (Nielsen-LeRoux et al. 2012; Richards and Goodrich-Blair 2009; Stilwell et al. 2018). Infective juvenile (IJ) nematodes carrying the bacteria actively seek out and infiltrate lepidopteran larvae and other hosts through cavities such as the mouth and anus (Akhurst 1980; Boemare, Akhurst, and Mourant 1993), where they release the bacteria, which then proliferate.

*Xenorhabdus nematophilus*, one of the most studied species in these genera, is an ideal microbe for the study of pathogen resource utilization due to its intimacy with its insect hosts’ environment. Lacking an external environmental phase limits the genetic adaptation that might occur were it to be exposed to environments highly variable in their resource availability (New et al. 2014). Moreover, nutrition plays a key role in the transition of the bacterial-nematode complex from mutualistic (in its nematode host) to pathogenic (in its insect host). Once in the haemolymph of the insect host, the *Steinernema* worms defaecate *X. nematophila,* which enters the exponential growth phase associated with increased virulence (Nielsen-LeRoux et al. 2012). So far there has been a limited number of studies investigating host diet effects on *X. nematophila* pathogenicity, and no studies directly investigating the outcome of the insect host internal-environment nutrient variation on this pathogen’s fitness (but see (Wilson et al. 2020)).

Holdbrook et al. (Holdbrook et al. 2024) characterized the host haemolymph nutrients of *S. littoralis* on 20 diets varying in their protein-to-carbohydrate ratio (Table S1). Cotter et al., (2019) showed, using the same 20 diets, that *S. littoralis* immune function is generally heightened in a high-protein environment, though the relationship between diet, immune gene expression and immunological effector molecules is complex. *Spodoptera* caterpillars can use dietary protein to reduce their susceptibility to *X. nematophila* (Wilson et al., 2020), demonstrating the potential for diet to impact its immune response (Cotter et al., 2019). It is also possible that dietary protein negatively impacts *X. nematophila* directly, via non-immune mechanisms (Wilson et al., 2020). To disentangle direct (nutritional) and indirect (host-mediated) effects on *Xenorhabdus* performance, we need to isolate the bacterium from its host and other potential microbes. One way of doing this is to characterise bacterial performance in synthetic haemolymphs (henceforth, *nutribloods*) that have similar nutritional properties to the natural haemolymphs generated when hosts feed on different diets but lacking the host’s immune effectors. Synthetic insect growth media have been around for many decades, mainly being used for *in vitro* cell cultures. For example, Grace’s insect medium was formulated to support growth of cells derived from the emperor gum moth *Opodiphthera* (formerly *Antheraea*) *eucalypti* and was itself modified from Wyatt’s medium, which was developed to mimic the haemolymph of the silkworm *Bombyx mori* (Schlaeger 1996). However, we are not aware of any previous studies that have synthesised multiple haemolymphs mimicking those of a single insect host-species feeding on different diets.

Here, we characterise the population growth characteristics of *X. nematophila* grown in nutritional environments (*nutribloods*) based on a nutritional analysis of *Spodoptera littoralis* haemolymph for larvae fed on 20 diets varying in their protein-to-carbohydrate ratios and concentrations (Holdbrook et al. 2024). The haemolymph nutrient variation was modelled statistically to produce 20 variable nutrient environments that the pathogen would experience based on those 20 host diets. *In vitro* pathogen growth was then measured over 30 hours in these 20 nutribloods. Based on *in vivo* findings (Wilson et al., 2020) and the fact that bacteria and virus-infected hosts survive better, and show a preference for protein-rich diets (Lee et al., 2006; Povey et al., 2009; Povey et al., 2014), we predicted that pathogen performance would be maximised in carbohydrate-rich, protein-poor environments.

## METHODS

### Bacterial cultures

Bacteria were originally supplied by the laboratory of Givaudan and colleagues (Montpellier University, France; *X. nematophila* F1D3 GFP labelled), maintained on nutrient agar at 4^°^C and stored in liquid culture at -80^°^C (1:1 nutrient broth culture: glycerol). To maintain virulence, bacteria were used to infect 6^th^ instar *S. littoralis* larvae. Harvested single colonies grown from haemolymph smeared NBTA agar plates (Sicard et al. 2004) were then grown in sterile nutrient broth for 24 h at 28^°^C shaking at 150 rpm. Stocks were made by mixing 500 µl of liquid culture with glycerol at a 1:1 ratio and stored again at -80^°^C. Prior to experiments, bacteria were revived from the frozen stores: 100 µl of frozen culture was added to 10 mL nutrient broth, which was then incubated for 16 h at 28^°^C shaking at 150 rpm.

### Synthetic haemolymph design

#### Nutrient sources

Except where stated, all nutrients used were obtained from Sigma Aldrich. The protein source used was bovine serum albumen (BSA, A7906), the carbohydrate source used was oyster glycogen (G8751), the lipid used was L-alpha-lecithin (Fisher Scientific 413102500) and all amino acids used were L-amino acids (A09416). The following sugars were used: Glucose (G8270), Fructose (F0127), Lactose (L3750), Sucrose (S9378), Trehalose (T9531).

#### Synthetic haemolymphs (nutribloods)

Haemolymph nutrients for larvae fed one of 20 diets (**Table S1**) were analysed through HPLC, uHPLC and lab nutrient assays (for details refer to (Holdbrook et al. 2024) and modelled using generalised additive models (**Table S2**). The model equations were then used to design the recipes for the 20 nutribloods (**Tables S3-5**). Mean values were used for nutrients that showed no statistical variation in larval haemolymph in response to diet, including carbohydrates and minerals. The five sugars, trehalose, glucose, fructose, lactose and sucrose were chosen to represent all sugars since these made up more than 95% of the haemolymph sugar content and the remaining simple sugars did not vary with diet (Holdbrook et al. 2024). Haemolymph amino acid ratios differed according to larval diet and hence were varied between nutribloods.

The starting point for all 20 nutribloods was based on the saline and vitamin content of Sigma Grace’s insect medium (SGIM, Sigma Aldrich, G8142). A basic salt solution (SGIM-saline) was prepared in sterile distilled water (**Table 1**: inorganic salts), the pH was adjusted to 4.5 with HCl and NaOH to dissolve the salts, and the solution was stored at 4^°^C.

**Table 1.** The results of the principal components analysis on the fourteen nutriblood variables. The variables with the greatest loadings for each PC (>0.3 or <-0.3) are shown in bold. (E) indicates that the nutrient is an essential amino acid.

| Nutrient | PC1 |  | PC2 |  |
| --- | --- | --- | --- | --- |
|  | Loading | Contribution | Loading | Contribution |
| Proteins | 0.270 | 7.273 | -0.226 | 5.122 |
| Carbohydrates | -0.124 | 1.529 | **0.427** | **18.258** |
| Glutamic acid | 0.156 | 2.442 | **-0.403** | **16.240** |
| Cysteine | 0.295 | 8.702 | 0.213 | 4.531 |
| Serine | 0.132 | 1.731 | **-0.421** | **17.723** |
| Arginine | 0.091 | 0.837 | **0.444** | **19.754** |
| Alanine | **0.304** | **9.227** | 0.187 | 3.509 |
| Aspartic acid (E) | **0.304** | **9.216** | 0.187 | 3.501 |
| Isoleucine (E) | **0.332** | **10.990** | -0.008 | 0.006 |
| Leucine (E) | **0.326** | **10.660** | -0.088 | 0.772 |
| Lysine (E) | **0.304** | **9.237** | 0.187 | 3.511 |
| Phenylalanine (E) | **0.330** | **10.868** | -0.048 | 0.227 |
| Tryptophan (E) | 0.284 | 8.061 | -0.183 | 3.337 |
| Valine (E) | **0.304** | **9.227** | 0.187 | 3.509 |
| Standard deviation | 3.004 |  | 2.157 |  |
| Variance explained (%) | 64.47 |  | 33.24 |  |
| Cumulative variance (%) | 64.47 |  | 97.72 |  |

The next stage was to make concentrated solutions of vitamins, sugars and amino acids: the SGIM-saline was filter-sterilized using 0.2 µM Corning sterile syringe filter and vitamins, sugars and amino acids were added from concentrated stock solutions to 10x the desired final concentrations (**Tables 3 and 4**). The solutions were made to a final volume of 10 mL by adding the stock solution to 9 mL of SGIM-saline solution, adjusting the pH to 6.4 and making up the final volume to 10 mL using SGIM-saline. These solutions were then filter-sterilized and divided into 1 mL aliquots in Eppendorf tubes and stored at -80^°^C.

To make the final nutriblood solutions, a 1 mL micronutrient solution was diluted to 9 mL with SGIM-saline, protein (BSA) and carbohydrate (glycogen) were added to the final concentration (**Table 5**). The pH was adjusted to 6.4 with HCl and NaOH to match the pH of lepidopteran haemolymph (Wyatt, 1961), and the final volume was made up to 9.8 mL with SGIM-saline. The solution was filter-sterilized with 0.2 µm cellulose acetate membrane filters (Sigma Aldrich, WHA69012502), and stored in 980 µL aliquots in Eppendorf tubes at -80^°^C. The volume was made up to 1 mL with the addition of 20 µL of a filter-sterilised 0.05 g/mL lipid stock solution once the tubes were retrieved from storage prior to use in assays.

#### Single-variable nutribloods (SVNs)

To isolate the effect of each nutrient group on bacterial growth, the other nutrient groups in solution were fixed at their mean value, whilst the nutrient of interest (protein, carbohydrate or amino acids) was increased in concentration. A set of 24 single-variable nutribloods (SVNs) were designed based on the variable nutrients in the 20 nutribloods. Each series included 6 levels, ranging from 0% of that nutrient to 100%. The 100% value was the maximum value plus 2 standard deviations from the nutriblood concentrations. This range was chosen as it covers the full range of variation a pathogen would experience in the nutrient environment in the host haemolymph.

### *In vitro* bacterial growth experiments

#### Preparation of bacterial cell culture for growth experiments

Bacteria were revived from frozen liquid stores after which time 2 mL was sub-cultured into 8 ml of nutrient broth and incubated for a further 4 h to reach log phase. The bacterial cells were washed to avoid the transfer of nutrients from nutrient broth into the growth media, following (Crawford et al., 2012). Briefly, 1 mL of sample was centrifuged for 6 min at 3000 g twice, removing the supernatant each time. Filter-sterilized SGIM-saline was used to re-suspend the cells in between the centrifugation steps. A 1 mL sample was subsequently used to produce a dilution series in SGIM-saline from which the total cell count was determined using a compound microscope (Zeiss Axioskop40) under fluorescence microscopy using a haemocytometer with improved Neubauer ruling. The remaining culture was diluted to 1 x 10^7^ cells/mL in the SGIM-saline solution, making the final starting concentration in each treatment 1 x 10^6^ cells/mL since 20 μL of bacterial cells were added to 180 μL of growth media.

#### Bacterial growth assays

Cell growth was determined at 28^°^C in Corning 96-well plates (Sigma Aldrich, CLS3595) using a SpectraMax Plus microtiter plate reader with SoftMax Pro software (Molecular Devices™). The wells of the plate contained 180 µL of one of the 20 nutribloods in quadruplets. SGIM-saline and SGIM were used as the negative and positive controls respectively. Twenty µL of bacterial culture was added to half of the wells (duplicates) and 20 µL of SGIM-saline was added to the other half as blank controls. The turbidity at 600 nm was determined every 10 min for 30 h and the plate was shaken for 30 s before each measurement. The experiment was repeated for three plates, with each plate containing all growth solutions and controls.

### Data analysis

Due to varying optical densities (OD) produced by lipids in the nutribloods, the minimum OD in each solution were variable. Prior to analysis, data were corrected by first subtracting a growth series from its corresponding blank series (to which no bacteria had been added). Then, the mean of the minimum 10 values in a series was subtracted from all the values in the series, producing a new zero-point. The adjustments removed the variation in OD caused by nutrients, maintaining only variation due to bacterial growth.

Data analysis was performed using the R statistical software (R version 4.3.0 ; R Core Team, 2014). To account for variation in haemolymph nutrient concentrations, data were standardized using the mean (µ) and standard deviation (σ), as per Cotter et al., (2011), allowing comparisons between nutrient groups; a standardized variable (Z) was produced from a nutrient variable (X) using the formula: (Z = (X-µ)/σ). Bacterial growth kinetics were calculated using the *Growthcurver* package v0.3.1 (Sprouffske, 2020). Initial analyses revealed that *in vitro K* (henceforth *max.OD* for consistency with (Wilson et al. 2020) was the metric most closely related to *in vivo* bacterial growth rate (Holdbrook 2019; Wilson et al. 2020) and so this is what we focus on here. This was then analysed using generalized additive models (GAM) in the *mgcv* package (v1.8.42 (Wood, 2011) and visualised via thin-plate spline plots created using the *fields* package (Nychka et al. 2017), following (Cotter et al., 2011). To complement the GAM analysis, the thin-plate regression splines were produced using the REML method for smoothing.

Because there was no significant variation in the haemolymph simple sugars, all haemolymph carbohydrates were analysed as an aggregate of glycogen and the sum of the simple sugars. Proteins and free amino acids were analysed separately. Due to the number of nutrients in the analysis, we also used principal components analysis (PCA) to reduce the number of dependent variables using the *prcomp* function in R.

Where appropriate, an information theoretic approach was taken for analysis (Whittingham et al., 2006). This allows the selection of multiple candidate models accounting for how much variation each explains based on the Akaike information criterion (AIC; (Burnham and Anderson 2004)). The AIC analysis was carried out using the *MuMIn* package (v1.47.5; (Bartoń 2023)) in R which, when combined with the *mgcv* package, ranks models based on the degrees of freedom used to create the smoothed curve. Where appropriate, model selection was carried out using evidence ratios, which provides the ratio of the model weights: an evidence ratio for the best model against model *X* is calculated by the Akaike weight of the best model/the Akaike weight of model *X*. The specific analyses varied, however they all included a ‘*Null* model’, which provided a baseline measure of variation without any nutritional information.

## RESULTS

### Covariation of *nutriblood* nutrients

The nutribloods contain multiple nutrients and many of these co-vary with each other (Figure 1). For example, levels of protein and carbohydrates in the nutribloods are strongly negatively correlated (r_s_ = -0.65) and the proportion of essential and non-essential amino acids are strongly positively correlated (r_s_ = 0.73). Whilst most individual amino acids are positively correlated with each other (r_s_ ≤ 1.00), some are significantly negatively correlated, such as arginine and alanine (r_s_ = -1.00) and isoleucine and tryptophan (r_s_ = -0.75). Due to such covariation, we started our analysis by conducting a principal components analysis.

**Figure 1.**
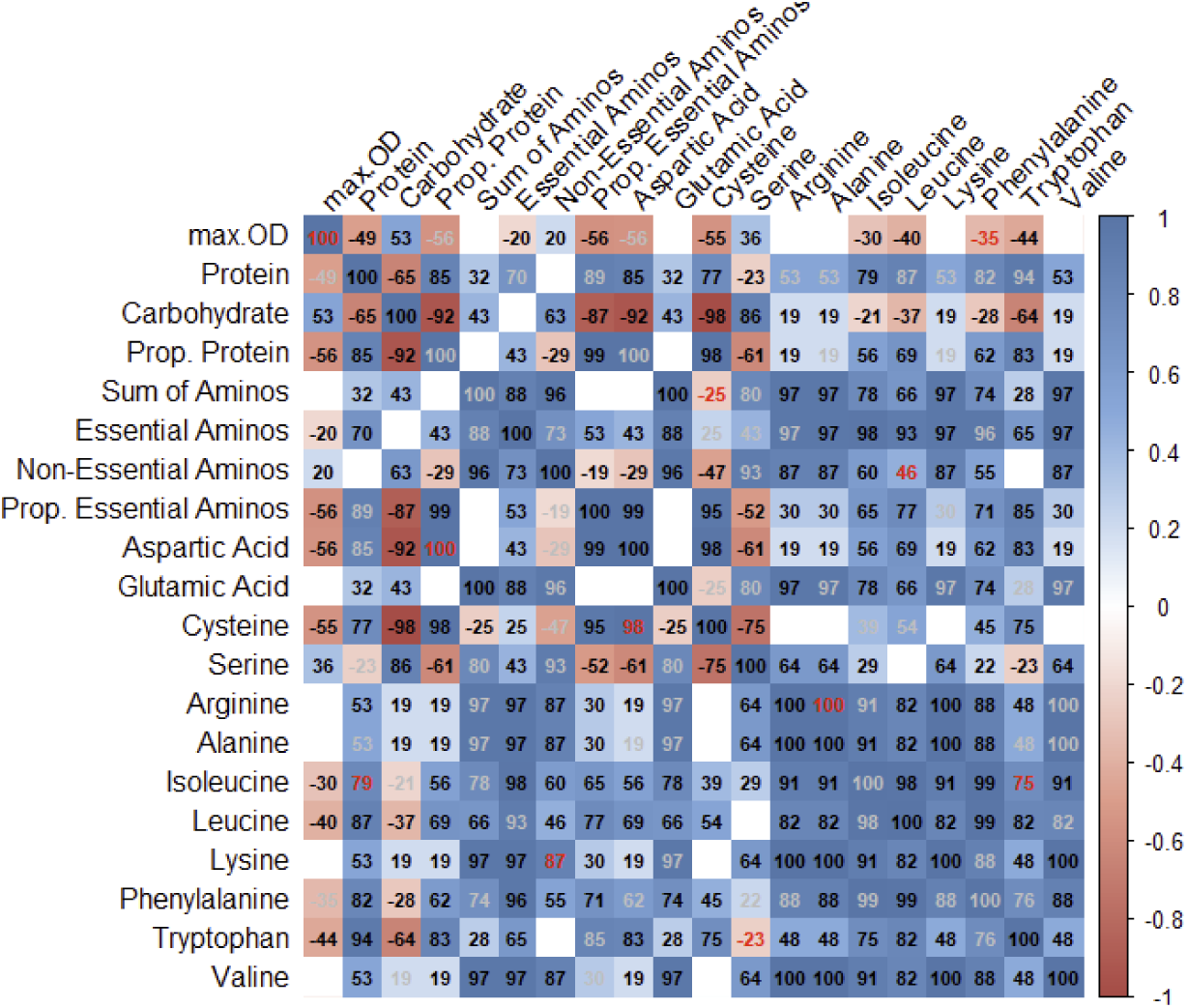
Spearman’s rank correlations between the variable nutrients in the nutribloods, expressed as percentages. Significant positive correlations are blue, significant negative correlations are red and non-significant correlations are white. The intensity of the colour indicates the strength of the correlation. The value of r*100 is indicated in each cell.

### Principal components analysis

A principal components analysis (PCA) that included all of the variable nutrients in the nutribloods revealed that the first two principal components explained nearly 98% of the variation in their nutritional properties (**Table 1**, **Figure 2**). The PCA loadings suggest that the PC1 axis is mostly positively associated with the concentration of protein and free amino acids in the nutribloods, especially the essential amino acids. In contrast, PC2 was mostly positively associated with the amount of carbohydrate in the nutribloods, as well the concentration of some non-essential amino acids, though it should be noted that the relationship between PC2 and both asparagine and cysteine was negative.

**Figure 2.**
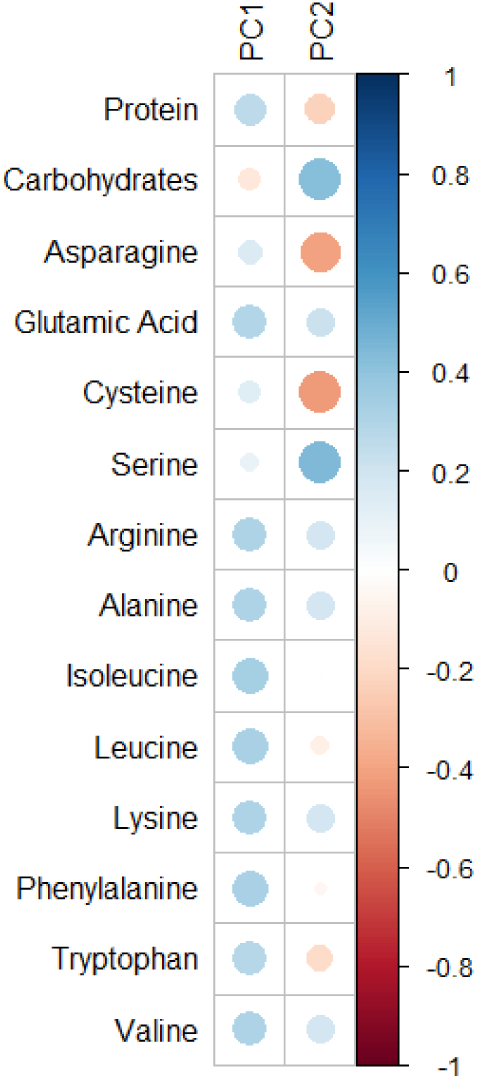
Principal components visualisation for the first two PCs. The size of the circles indicates the strength of the correlation and the colour indicates the direction (blue = positive, red = negative).

The relationship between these two principal component axes and bacterial growth in the 20 nutribloods was explored using GAMs. The top model (m4) included a significant interaction between the two principal components (**Table 2**). Together, PC1 and PC2 explained around 53% of the variation in bacterial growth, with PC1 being associated with a non-linear relationship with *max.OD*, peaking at relatively low values, and PC2 being positively correlated with *max.OD* (**Figure 3a**). This indicates that bacterial growth is highest when levels of protein and the majority of amino acids are low (low levels of PC1) and when carbohydrates and serine are high, and asparagine and cysteine are low (high levels of PC2)

**Figure 3.**
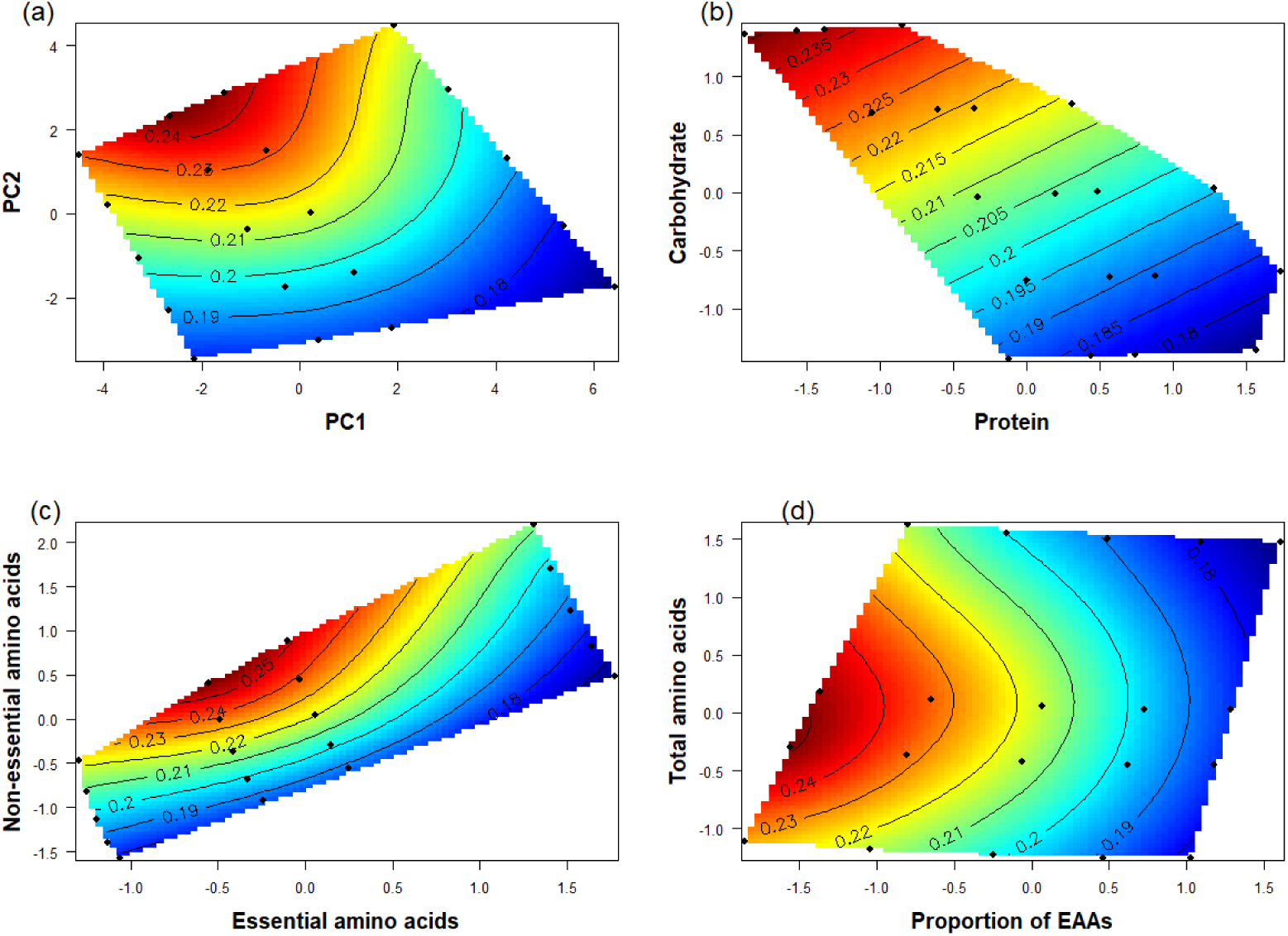
Effect of nutrients on bacterial growth. 2D contour plots compare a dependent variable (*max.OD*) on the z-axis with two nutritional attributes of the nutribloods simultaneously. Contour lines represent the type of relationship between the dependent variable and the two independent variables. Colour represents the strength of effect, with blue indicating the weakest effect going up to red which represents the strongest effect. Nutrient concentrations were standardized for analysis to allow comparison. a) PC2 vs PC1; (b) carbohydrate vs protein; (c) non-essential vs essential amino acids; (d) total amino acids vs the proportion of those amino acids that are essential.

**Table 2.** Summary tables explaining variation in bacterial growth (*max.OD*) in relation to the nutrient composition of the nutribloods as reflected in the first two principal components. (a) AIC comparison table for models containing PC1 and PC2 representation of nutrients in the nutribloods as explanatory variables. (b) Summary information for each model in the AIC comparison table.

**(a) Model AICs: Nutriblood PC1 and PC2**
| Model | df | k | AICc | delta | weight | R<sup>2</sup> |
| --- | --- | --- | --- | --- | --- | --- |
| m4. PC1 * PC2 | 8 | 3 | -525.9 | 0.00 | 1.00 | 0.533 |
| m3. PC1 + PC2 | 8 | 2 | -510.8 | 15.07 | 0.00 | 0.468 |
| m2. PC2 | 4 | 1 | -477.4 | 48.49 | 0.00 | 0.242 |
| m1. PC1 | 4 | 1 | -467.4 | 58.50 | 0.00 | 0.176 |
| m0. Null | 2 | 0 | -448.9 | 76.99 | 0.00 | 0.000 |

**(b) Nutriblood PC1 and PC2 model summaries**
| Model | Terms | edf | F-value | P-value |
| --- | --- | --- | --- | --- |
| m1 | PC1 | 1.953 | 5.501 | <0.001 |
| m2 | PC2 | 1.995 | 8.778 | <0.001 |
| m3 | PC1 + | 3.354 | 12.41 | <0.001 |
|  | PC2 | 2.054 | 14.68 | <0.001 |
| m4 | PC1 + | 2.679 | 11.33 | <0.001 |
|  | PC2 + | 1.860 | 6.21 | <0.001 |
|  | PC1 : PC2 | 0.961 | 12.46 | <0.001 |

### Macronutrients and *in vitro* bacterial growth

To explore these patterns further, we analysed a number of the specific components of PC1 and PC2, starting with the two caloric macronutrients. Bacterial growth declined with increasing nutriblood protein and increased with increasing nutriblood carbohydrate, with *max.OD* peaking in nutrient space that was low in protein and high in carbohydrate (**Table 3a,b; Figure 3b**). Together, these two macronutrients explained ∼31% of the variation in bacterial growth.

**Table 3.** Summary tables explaining variation in bacterial growth (max.OD) in relation to the macronutrient composition of the nutribloods. (a) AIC comparison table for models containing macronutrients in the nutribloods as explanatory variables. (b) Summary information for each model in the AIC comparison table. (c) Summary information from models testing the effect of proportion of dietary protein (relative to all macronutrients) on *max.OD*.

**(a) Model AICs: Nutriblood protein and carbohydrate**
| Model | df | k | AICc | delta | weight | R<sup>2</sup> |
| --- | --- | --- | --- | --- | --- | --- |
| m3. Protein + Carbs | 3 | 2 | -490.7 | 0.00 | 0.420 | 0.308 |
| m4. Protein * Carbs | 3 | 3 | -490.6 | 0.00 | 0.420 | 0.308 |
| m2. Carbs | 3 | 1 | -488.6 | 2.03 | 0.152 | 0.288 |
| m1. Protein | 2 | 1 | -482.8 | 7.82 | 0.008 | 0.253 |
| m0. Null | 2 | 0 | -448.9 | 41.74 | 0.000 | 0.000 |

**(b) Nutriblood protein and carbohydrate model summaries**
| Model | Terms | edf | F-value | P-value |
| --- | --- | --- | --- | --- |
| m1 | Protein | 0.976 | 10.086 | <0.001 |
| m2 | Carbs | 0.980 | 12.056 | <0.001 |
| m3 | Protein | 0.806 | 1.039 | 0.022 |
|  | Carbs | 0.927 | 3.170 | <0.001 |
| m4 | Protein | 0.806 | 1.039 | 0.022 |
|  | Carbs | 0.927 | 3.169 | <0.001 |
|  | Protein: Carbs | 0.001 | 0.000 | 1.000 |

**(c) Proportion of nutriblood protein model summaries**
| Nutrient | R<sup>2</sup> | edf | F-value | P-value |
| --- | --- | --- | --- | --- |
| Proportion Protein | 0.306 | 1.861 | 13.119 | <0.001 |

It is worth noting, however, that the amounts of protein and carbohydrate in the nutribloods are strongly negatively correlated with each other (Pearson’s r = -0.751, df = 118, P < 0.001) and that the relative *proportions* of the two macronutrients in the nutriblood explained a similar amount of variation in bacterial growth as the top model based on absolute quantities of the two macronutrients (R^2^ = 0.306; **Table 3c**).

### Free amino acids and *in vitro* bacterial growth

In the PCA analysis, it appeared that PC1 was mostly associated with the essential amino acids (EEA), whereas PC2 was mostly associated with non-essential amino acids (non-EAA). To explore this further, we first looked at the apparent contrasting effects of essential and non-essential amino acids on bacterial growth. The top model (m4, **Table 4a**) was one in which there was a significant interaction between these two main effects, but the simpler additive effects model (m3) explained a similar amount of variation (m4 R^2^ = 0.551, m3 R^2^ = 0.546; **Table 4b**). Bacterial growth peaked in a nutrient space that contained relatively low concentrations of EAA and medium to high concentrations of non-EAA (**Figure 3c**). Reflecting these findings, there was a decline in *max.OD* as the proportion of EAA relative to all amino acids increased and bacterial growth actually peaked at intermediate levels of all free amino acids (**Table 5**, **Figure 3d**).

**Table 4.** Summary tables explaining variation in bacterial growth (max.OD) in relation to the amounts of essential and non-essential amino acids of the nutribloods. (a) AIC comparison table for models containing essential and non-essential in the nutribloods as explanatory variables. (b) Summary information for each model in the AIC comparison table.

**(a) Model AICs: EAA and non-EAA**
| Model | df | k | AICc | delta | weight | R<sup>2</sup> |
| --- | --- | --- | --- | --- | --- | --- |
| m4. EAA * Non-EAA | 8 | 3 | -536.2 | 0.00 | 0.631 | 0.551 |
| m3. EAA + Non-EAA | 8 | 2 | -535.1 | 1.08 | 0.369 | 0.546 |
| m1. EAA | 4 | 1 | -472.4 | 63.79 | 0.000 | 0.201 |
| m2. Non-EAA | 4 | 1 | -471.4 | 64.78 | 0.000 | 0.195 |
| m0. Null | 2 | 0 | -448.9 | 87.25 | 0.000 | 0.000 |

**(b) EAA and non-EAA model summaries**
| Model | Terms | edf | F-value | P-value |
| --- | --- | --- | --- | --- |
| m1 | EAA | 2.543 | 7.468 | <0.001 |
| m2 | Non-EAA | 2.737 | 7.187 | <0.001 |
| m3 | EAA + | 2.525 | 23.34 | <0.001 |
|  | Non-EAA | 3.220 | 22.92 | <0.001 |
| m4 | EAA + | 2.214 | 17.844 | <0.001 |
|  | Non-EAA + | 2.812 | 9.965 | <0.001 |
|  | EAA : Non-EAA | 0.827 | 0.752 | 0.003 |

**Table 5.** Summary tables explaining variation in bacterial growth (*max.OD*) in relation to the total amount of free amino acids and proportion of essential amino acids of the nutribloods. (a) AIC comparison table for models containing essential and non-essential amino acids in the nutribloods as explanatory variables. (b) Summary information for each model in the AIC comparison table.

**(a) Model AICs: All free amino acids and proportion EAA**
| Model | df | k | AICc | delta | weight | R<sup>2</sup> |
| --- | --- | --- | --- | --- | --- | --- |
| m4. sum.aminos * prop.EAA | 5 | 3 | -520.0 | 0.00 | 0.5 | 0.465 |
| m3. sum.aminos +<br>prop.EAA | 5 | 2 | -520.0 | 0.00 | 0.5 | 0.465 |
| m2. prop.EAA | 3 | 1 | -495.0 | 25.02 | 0.0 | 0.325 |
| m1. sum.aminos | 4 | 1 | -468.8 | 51.25 | 0.0 | 0.171 |
| m0. Null | 2 | 0 | -448.9 | 71.11 | 0.0 | 0.000 |

**(b) All free amino acids and proportion EAA**
| Model | Terms | edf | F-value | P-value |
| --- | --- | --- | --- | --- |
| m1 | sum.aminos | 2.016 | 6.157 | <0.001 |
| m2 | prop.EAA | 0.983 | 15.33 | <0.001 |
| m3 | sum.aminos | 2.108 | 7.727 | <0.001 |
|  | prop.EAA | 0.985 | 16.079 | <0.001 |
| m4 | sum.aminos | 2.108 | 7.727 | <0.001 |
|  | prop.EAA | 0.985 | 16.079 | <0.001 |
|  | sum.aminos |  |  |  |
|  | prop.EAA | 0.000 | 0.000 | 1.000 |

As there is strong covariation between most of the individual amino acids in the nutribloods (Figure 1), we did not analyse the effects of specific amino acids further.

### Single variable nutribloods (SVN) and *in vitro* bacterial growth

In order to explore some of these patterns in a more controlled environment, we developed another series of 24 synthetic *single variable nutribloods* (SVNs) in which all nutrients were fixed at an average level but then concentrations of the three main nutrient groups were varied systematically around their means in the nutribloods ± 2 standard deviations and *in vitro* bacterial growth was again quantified using *max.OD*. This revealed that bacterial growth declined non-linearly with protein concentration (R^2^ = 0.783) and increased linearly with increasing carbohydrate (R^2^ = 0.851), mirroring what we observed with the 20 nutribloods (Figure 3b). In contrast, bacterial growth rate in the SVNs increased non-linearly with the concentration of free amino acids (R^2^ = 0.648), with *max.OD* being maximised in synthetic haemolymphs with relatively high carbohydrate and amino acid concentrations and relatively low protein concentrations (Table 6, Figure 4).

**Figure 4.**
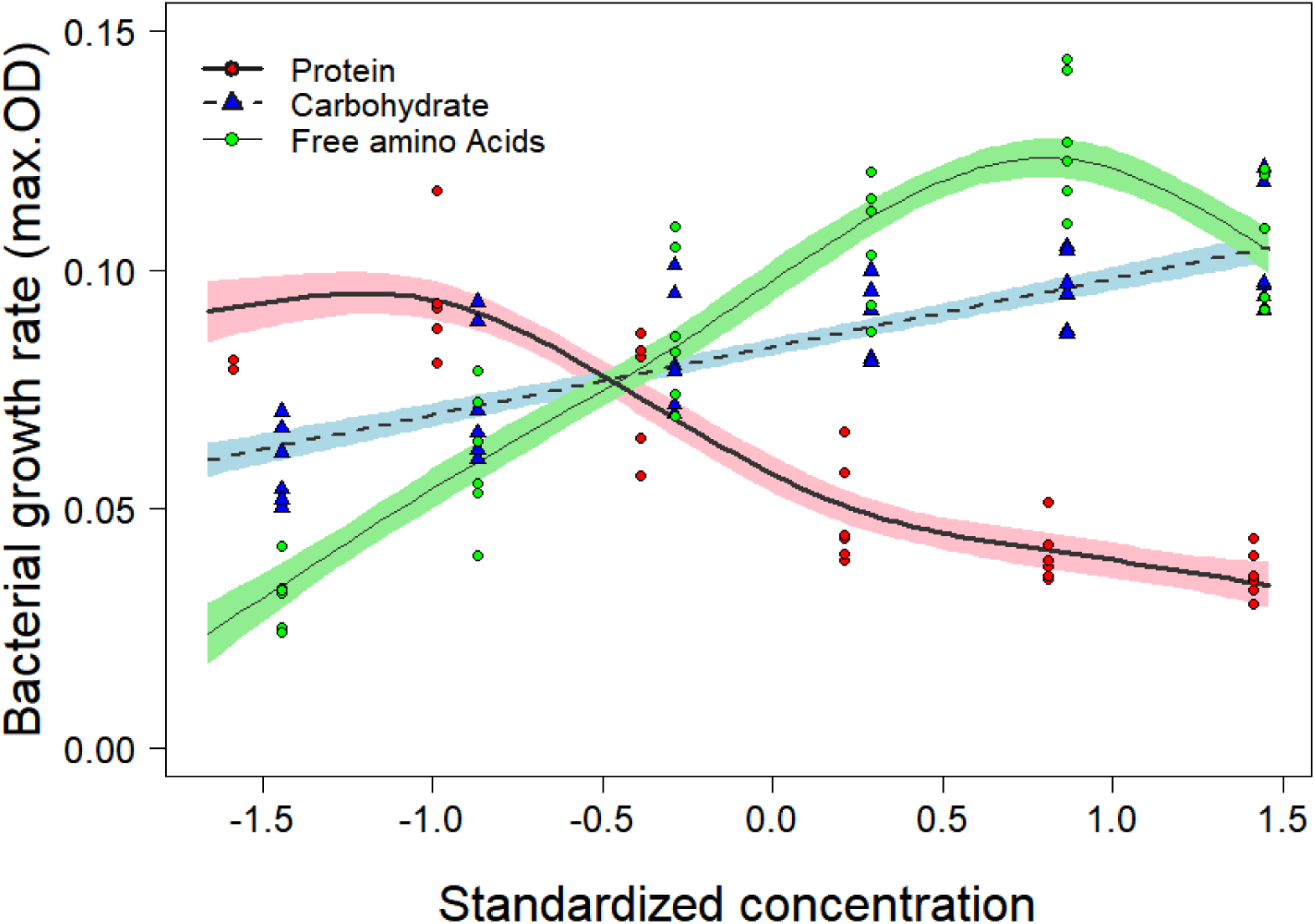
Effect of three key nutrient groups on bacterial growth. The lines and shading are the fitted values and standard errors from the GAM models for each of protein, carbohydrate and amino acids; symbols are the individual point. Note that two outliers were excluded which had a *max.OD* of zero.

**Table 6.**
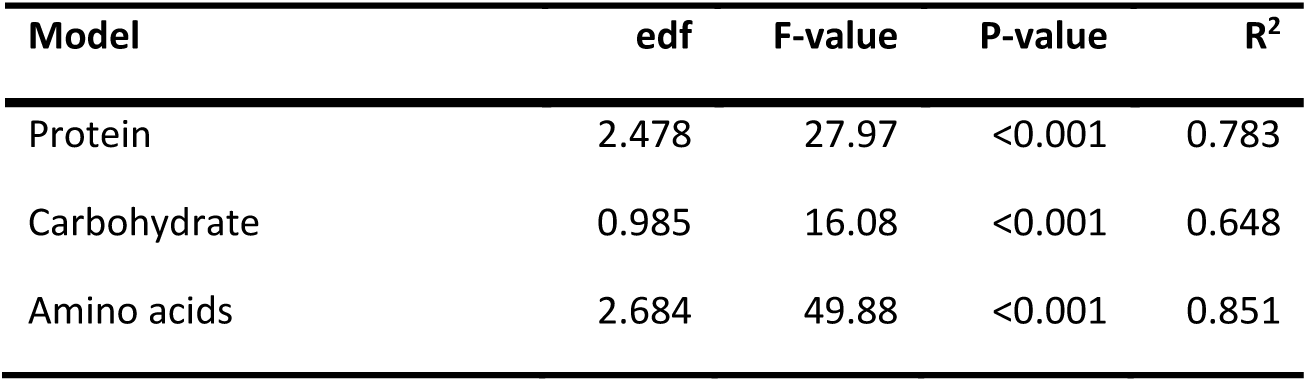
Summary table explaining variation in bacterial growth (max.OD) in relation to variation in three key nutrient groups using Single Variable Nutribloods (SVNs). Summary information for each model in the AIC comparison table.

## DISCUSSION

In a previous study, we established how macronutrient variation in twenty chemically-defined diets (Cotter et al., 2011) manifests itself in variation in the nutritional profiles of *Spodoptera littoralis* haemolymph (Cotter et al. 2011; Holdbrook et al. 2024). In the present study, we used these data to design twenty ‘*nutribloods*’ that reflect the nutritional properties of the haemolymphs of insects that had fed on these twenty diets. The rationale for this was to establish how the growth of entomopathogenic bacteria varies in relation to the nutritional properties of the haemolymph in the absence of its insect host’s immune system. More generally, we provide recipes for a range of nutritional media that reflect the haemolymph of insects that can be used to compare the performance of a range of entomopathogens both in isolation and in combination. The model pathogen used in this system was *Xenorhabdus nematophila*, a gram-negative bacterium that is adapted to growing in insect blood. Our results revealed that *in vitro* bacterial growth tended to be highest in nutribloods that were carbohydrate-rich and protein-poor, which is consistent with *in vivo* studies (Wilson et al., 2020). However, the levels of these two macronutrients in the nutribloods (and in real haemolymphs) strongly covary with levels of free amino acids (Figure 1, Table 3) and further analyses suggest that bacterial growth increases with the levels of free amino acids when macronutrients are at intermediate levels.

### Macronutrient effects

A previous study (Wilson et al., 2020), using a more limited set of (six) chemically-defined diets suggested that the outcome of the parasitic interaction between *S. littoralis* and *X. nematophila* is associated with the amount of protein in the host diet. This is consistent with earlier host-pathogen studies using *Spodoptera* caterpillars showing that dietary protein tends to improve host performance in the face of pathogenic challenges, whilst dietary carbohydrates tends to reduce it (Lee et al., 2006; Povey et al., 2009; Povey et al., 2014). *Spodoptera* haemolymph properties are tightly regulated in comparison to the diets they eat, whilst blood protein tends to increase with dietary protein (Cotter et al., 2011; Povey et al., 2009), carbohydrate levels are more strongly regulated (Povey et al. 2009; Cotter et al. 2011; Holdbrook et al. 2024). Indeed, the levels of simple sugars (glucose, trehalose, etc.) did not vary across our twenty nutribloods, reflecting what is observed in host haemolymphs (Holdbrook et al. 2024). In the present study, we found that *in vitro* bacterial growth was positively associated with the amount of carbohydrate in the nutriblood and negatively associated with the amount of protein (Figure 1, Figure 3b). However, due to post-ingestive nutrient regulation by the insect, levels of the two macronutrients in the haemolymph (and, by extension, in the nutribloods) are strongly negatively correlated (r = -0.751, P < 0.001; Figure 3b), making it difficult to distinguish between positive carbohydrate effects on bacterial growth versus negative protein effects, or indeed other correlated nutrient effects (but see (Wilson et al., 2020)). However, the single variable nutribloods fixed most components of the nutribloods at intermediate levels and varied carbohydrates and proteins independently. These confirmed the negative effects of protein on bacterial growth and the independent positive effects of carbohydrates.

### Amino acid effects

In contrast to the invariant levels of simple sugars, free amino acids did vary significantly across nutribloods, with bacterial growth peaking at intermediate levels (Figure 3d). However, when protein and carbohydrate levels were fixed at intermediate levels and the concentration of free amino acids was varied (in SVNs), bacterial growth increased non-linearly with increasing amino acid levels (Figure 4), suggesting that they are using these as a food source, but that excess free amino acids may become toxic, consistent with Bertrand’s rule (Raubenheimer, Lee, and Simpson 2005). The difference between these two experiments is likely due to the fact that, in the twenty nutribloods, protein and amino acid levels tend to strongly covary (*r* = 0.493, P< 0.001), and so nutribloods rich in amino acids, which can be used for growth, are also rich in proteins, which are detrimental to bacterial growth.

The nutribloods appear to provide some evidence for contrasting effects of essential (EAA) and non-essential (non-EAA) amino acids, with bacterial growth being inhibited as the proportion of EAA in the free amino acid pool increases (Figure 3c, 3d). However, this again is likely to be an artefact of the fact that EAA levels in the nutribloods are positively correlated with levels of (damaging) proteins (*r* = 0.630, P< 0.001), whereas the concentration of non-EAA tended to be positively correlated with levels of (favourable) carbohydrates (*r* = 0.523, P< 0.001). Thus, the proportion of EAA in the amino acid pool probably reflects the proportion of protein in the macronutrient pool. In the future, single variable nutribloods could be used to test this hypothesis directly.

### Protein may interfere with bacterial osmoregulation

Abisgold & Simpson (1987) found that increasing protein concentration in the diet of *Locusta migratoria* increased the concentration of amino acids in the haemolymph and haemolymph osmolality. Using *Spodoptera littoralis*, Wilson et al., (2020) found the same effects using 6 diets that varied independently in their protein and carbohydrate content; high-protein diets increased blood osmolality and high osmolality decreased bacterial growth rates *in vivo* and *in vitro*. The ability of a cell to adapt to changes in their osmotic environment, i.e., osmoregulation, is important for the maintenance of turgor pressure across the cellular membrane (Kempf and Bremer 1998; Csonka and Hanson 1991). This, in turn, determines the cell’s ability to counteract osmotic stress and therefore its capacity to proliferate (Csonka 1989; Tempest, Meers, and Brown 1970). The findings of Wilson et al., (2020) highlight changes in osmolality as a possible mechanism for the observed negative ‘protein effect’ on bacterial population growth. Since the feeding interval of insects is decided by haemolymph osmolality (Simpson and Abisgold, 1985), this must be very tightly regulated. A host-dependent entomopathogen like *X. nematophila* would be sensitive to osmotic stress, due to fairly limited variability in its osmotic environment. The reduction in bacterial population growth in response to mid-to-high protein levels and high amino acid levels may be partially due to cells changing state upon sensing a high-osmolality environment (Cesar et al. 2020).

Prokaryotic organisms depend on the uptake of free amino acids for osmoregulation (Kempf and Bremer 1998; Krell et al. 2010; Tempest, Meers, and Brown 1970). This means that after amino acids from a host meal raise haemolymph osmolality, the pathogen absorbs these very same amino acids in reaction to the osmotic changes. This would cause a reduction in the haemolymph amino acid concentration signalling to the host to seek out more protein. Consistent with this, Cotter et al., (2011) failed to observe a change in host protein diet-choice using an immune elicitor, unlike previous experiments that used live pathogens (Lee et al., 2006; Povey et al., 2009). Investigating the effect of amino acids rather than whole proteins may provide clues as to how protein may be altering the reaction of *X. nematophila* to its osmotic environment.

### Further uses for nutribloods

A key motivation for the development of these nutribloods was to better understand the mechanisms by which nutrition impacts an insect’s capacity to cope with entomopathogens by disentangling insect-mediated factors on pathogen fitness (e.g. the host’s immune response) from the direct impacts of host nutrition (e.g. mediated by the bacteria’s nutritional requirements). In a follow up paper (Holdbrook et al., in prep), we combine the *in vitro* bacterial growth data generated using these nutribloods with *in vivo* data generated by sampling the bacterium directly from the host itself to determine the discrepancy between how the bacterium grows in the presence and absence of the host’s immune system and other host-mediated effects.

Future studies can use these nutribloods to compare how host nutrition affects different entomopathogens; not just other bacteria but other extracellular parasites of the blood system including entomophagous fungi, protists, nematodes, etc. For example, one potential use of this system is to elucidate the nutritional requirements of the nematode symbionts of *X. nematophila*, such as *S. carpocapsae*, which are currently unknown (Richards and Goodrich-Blair 2009). There is also scope for using these nutribloods to explore the effect of host nutrition on competition between entomopathogens during co-infections. Finally, nutribloods could be used to explore the role of host insects on extracellular symbionts.

## Conclusion

To our knowledge, this study is the first to systematically develop a range of synthetic haemolymphs (nutribloods) that reflect the nutritional properties of haemolymphs when the insect (*S. littoralis*) feeds on a range of diets varying in their macronutrient compositions. We then tested the impact of these different nutribloods on the growth of an entomopathogen (*X. nematophila*). We provide strong evidence that the growth rate of this pathogen is greatest in a high-carbohydrate, low-protein nutrient space. We also studied bacterial growth in a less constrained nutriblood system in which most nutrients were maintained at intermediate levels but one of the three main nutrient groups were systematically varied. Whilst nutriblood protein constrains *X. nematophila* growth, both carbohydrate and free amino acids facilitate it, suggesting that these are both used as a nutrient source by the bacteria. In addition, we have provided an experimental framework for testing the role of nutrition in host-entomopathogen interactions, as well as for studying nutritional effects on entomopathogen co-infections and microbial symbionts.

