## Supplementary material for "Nutribloods: novel synthetic lepidopteran haemolymphs for understanding insect-microbe interactions *in vitro*"

**SUPPLEMENTARY TABLES**

**Table S1:** Twenty diets fed to *Spodoptera littoralis* caterpillars to produce varying haemolymph nutritional environments on which the synthetic haemolymphs were based.

| Diet.no | P:C | P:C ratio | conc | ratio prot | %prot | %carb |
|---|---|---|---|---|---|---|
| 1 | 10.5 : 52.5 | 1:5 | 63 | 0.17 | 10.5 | 52.5 |
| 2 | 7:35 | 1:5 | 42 | 0.17 | 7 | 35 |
| 3 | 5.6 : 28 | 1:5 | 33.6 | 0.17 | 5.6 | 28 |
| 4 | 2.8 :14 | 1:5 | 16.8 | 0.17 | 2.8 | 14 |
| 5 | 21:42 | 1:2 | 63 | 0.33 | 21 | 42 |
| 6 | 14:28 | 1:2 | 42 | 0.33 | 14 | 28 |
| 7 | 11.2 : 22.4 | 1:2 | 33.6 | 0.33 | 11.2 | 22.4 |
| 8 | 5.6 : 11.2 | 1:2 | 16.8 | 0.33 | 5.6 | 11.2 |
| 9 | 31.5 : 31.5 | 1:1 | 63 | 0.50 | 31.5 | 31.5 |
| 10 | 21:21 | 1:1 | 42 | 0.50 | 21 | 21 |
| 11 | 16.8 : 16.8 | 1:1 | 33.6 | 0.50 | 16.8 | 16.8 |
| 12 | 8.4 : 8.4 | 1:1 | 16.8 | 0.50 | 8.4 | 8.4 |
| 13 | 42 : 21 | 2:1 | 63 | 0.67 | 42 | 21 |
| 14 | 28:14 | 2:1 | 42 | 0.67 | 28 | 14 |
| 15 | 22.4 : 11.2 | 2:1 | 33.6 | 0.67 | 22.4 | 11.2 |
| 16 | 11.2 : 5.6 | 2:1 | 16.8 | 0.67 | 11.2 | 5.6 |
| 17 | 52.5 : 10.5 | 5:1 | 63 | 0.83 | 52.5 | 10.5 |
| 18 | 35:7 | 5:1 | 42 | 0.83 | 35 | 7 |
| 19 | 28 : 5.6 | 5:1 | 33.6 | 0.83 | 28 | 5.6 |
| 20 | 14 : 2.8 | 5:1 | 16.8 | 0.83 | 14 | 2.8 |

**Table S2:** Selected models for each haemolymph nutritional component used in Nutriblood predictions

| | Nutrients | Best model | Deviance explained | Transformation | Best model - transformed | Deviance explained |
|---|---|---|---|---|---|---|
| AMINO ACIDS | Leucine (LEU) | P | 26.6% | X^0.275 | P*C | 29.2% |
| | Tryptophan (TRP) | P | 25% | X^0.325 | P | 20.1% |
| | Aspartate (ASP) | P*C | 16.7% | X^0.45 | P*C | 18.2% |
| | Phenylalanine (PHE) | P | 16.7% | X^0.35 | P | 17.7% |
| | Cysteine (CYS) | P | 15.7% | Log(X) | P | 4.7% |
| | Isoleucine (ILE) | P+C | 14.4% | X^0.125 | P | 15.7% |
| | Serine (SER) | C | 13.5% | X^0.5 | C | 13.8% |
| | Glutamine (GLN) | C | 9.6% | X^0.3 | C | 10.7% |
| | Methionine (MET) | P | 8.9% | X^0.575 | P | 8.5% |
| | GABA | P | 7.9% | X^0.375 | Int | 0% |
| | Glycine (GLY) | P | 7.7% | X^0.55 | Int | 6.4% |
| | Threonine (THR) | C | 7.6% | X^0.225 | C | 5.9% |
| | Valine (VAL) | C | 6.3% | X^0.075 | P | 7.5% |
| | Glutamate (GLU) | C | 6.2% | X^0.375 | C | 5.9% |
| | Histadine (HIS) | P | 4.4% | X^0.425 | Int | 0% |
| | Asparagine (ASN) | Int | 0% | X^0.275 | Int | 0% |
| | Arginine (ARG) | Int | 0% | -1*X^-0.25 | Int | 0% |
| | Alanine (ALA) | Int | 0% | X^0.6 | Int | 0% |
| | Tyrosine (TYR) | Int | 0% | X^0.275 | Int | 0% |
| | Lysine (LYS) | Int | 0% | -1*X^-0.1 | P | 9.2% |
| | Proline | Int | 0% | X^0.45 | Int | 0% |
| SIMPLE SUGARS | Glucose | Int | 0% | X^0.225 | Int | 0% |
| | Fructose | Int | 0% | X^0.225 | Int | 0% |
| | Lactose | Int | 0% | X^0.225 | Int | 0% |
| | Sucrose | Int | 0% | X^0.2 | Int | 0% |
| | Sorbitol | Int | 0% | X^0.2 | Int | 0% |
| | Stachyose | P | 6.3% | X^0.225 | P | 7.1% |
| | Trehalose | Int | 0% | X^0.2 | P | 6.8% |
| MACRO-NUTR. | Carbohydrate | P+C | 27.9% | X^0.475 | P+C | 36.5% |
| | Protein | P | 36.4% | -1*X^-0.675 | P | 49.8% |
| | Lipid | Int | 0% | X^0.35 | Int | 0% |

**Table S3.** Concentrations of nutrients that were added in the same amounts to all of the nutribloods.

| | Nutrient | Concentration (g/L) |
|---|---|---|
| INORGANIC SALTS | $CaCl_2$ | 1.00E+00 |
| | KCl | 2.24E+00 |
| | $MgCl_2$ | 1.07E+00 |
| | $MgSO_4$ | 1.36E+00 |
| | $NaHCO_3$ | 3.50E-01 |
| | $Na_2HPO_4$ | 8.76E-01 |
| VITAMINS | Aminobenzoic acid | 2.00E-05 |
| | Biotin | 1.00E-05 |
| | Choline chloride | 2.00E-04 |
| | Folic acid | 2.00E-05 |
| | myo-inositol | 2.00E-05 |
| | Nicotinic acid | 2.00E-05 |
| | Pantothenic acid | 2.00E-05 |
| | Pyridoxine | 2.00E-05 |
| | Riboflavin | 2.00E-05 |
| | Thiamine | 2.00E-05 |
| SUGARS | Glucose | 9.65E-02 |
| | Fructose | 1.59E-02 |
| | Lactose | 2.40E-02 |
| | Sucrose | 2.62E-03 |
| | Trehalose | 3.22E-02 |
| AMINO ACIDS | Asparagine | 1.79E-06 |
| | Glutamine | 2.31E-05 |
| | Glycine | 3.61E-03 |
| | Histidine* | 2.36E-02 |
| | Methionine* | 4.72E-04 |
| | Proline | 6.43E-04 |
| | Threonine* | 1.71E-03 |
| | Tyrosine | 5.66E-04 |

**Table S4.** Concentrations (g/L) of the amino acids that were added in variable amounts to the nutribloods.

| Nutriblood | Aspartic acid | Glutamic acid | Serine | Alanine | Cysteine | Arginine * | Valine* | Tryptophan * | Phenylalanine * | Isoleucine * | Leucine* | Lysine* |
|---|---|---|---|---|---|---|---|---|---|---|---|---|
| 1 | 1.81E-04 | 7.11E-04 | 2.92E-02 | 1.40E-02 | 3.02E-03 | 8.17E-03 | 7.40E-03 | 3.74E-08 | 2.25E-03 | 2.42E-03 | 6.72E-03 | 7.16E-02 |
| 2 | 1.72E-04 | 5.28E-04 | 2.27E-02 | 1.06E-02 | 3.02E-03 | 5.51E-03 | 5.02E-03 | 4.80E-08 | 1.41E-03 | 1.48E-03 | 3.95E-03 | 5.37E-02 |
| 3 | 1.68E-04 | 4.66E-04 | 2.04E-02 | 9.37E-03 | 3.02E-03 | 4.70E-03 | 4.30E-03 | 5.22E-08 | 1.17E-03 | 1.21E-03 | 3.04E-03 | 4.78E-02 |
| 4 | 1.61E-04 | 3.59E-04 | 1.61E-02 | 7.13E-03 | 3.02E-03 | 3.41E-03 | 3.14E-03 | 6.07E-08 | 7.82E-04 | 7.93E-04 | 1.57E-03 | 3.80E-02 |
| 5 | 2.47E-04 | 7.04E-04 | 2.49E-02 | 1.40E-02 | 3.41E-03 | 8.17E-03 | 7.40E-03 | 3.93E-07 | 2.60E-03 | 2.73E-03 | 7.96E-03 | 7.16E-02 |
| 6 | 2.36E-04 | 5.23E-04 | 1.90E-02 | 1.06E-02 | 3.41E-03 | 5.51E-03 | 5.02E-03 | 2.50E-07 | 1.64E-03 | 1.67E-03 | 4.92E-03 | 5.37E-02 |
| 7 | 2.31E-04 | 4.61E-04 | 1.69E-02 | 9.37E-03 | 3.41E-03 | 4.70E-03 | 4.30E-03 | 1.93E-07 | 1.35E-03 | 1.37E-03 | 3.90E-03 | 4.78E-02 |
| 8 | 2.23E-04 | 3.55E-04 | 1.30E-02 | 7.13E-03 | 3.41E-03 | 3.41E-03 | 3.14E-03 | 7.97E-08 | 9.16E-04 | 9.06E-04 | 2.21E-03 | 3.80E-02 |
| 9 | 3.32E-04 | 6.97E-04 | 2.08E-02 | 1.40E-02 | 3.88E-03 | 8.17E-03 | 7.40E-03 | 7.70E-07 | 3.01E-03 | 3.11E-03 | 9.38E-03 | 7.16E-02 |
| 10 | 3.18E-04 | 5.17E-04 | 1.54E-02 | 1.06E-02 | 3.88E-03 | 5.51E-03 | 5.02E-03 | 4.65E-07 | 1.91E-03 | 1.91E-03 | 6.05E-03 | 5.37E-02 |
| 11 | 3.13E-04 | 4.56E-04 | 1.35E-02 | 9.37E-03 | 3.88E-03 | 4.70E-03 | 4.30E-03 | 3.43E-07 | 1.58E-03 | 1.56E-03 | 4.92E-03 | 4.78E-02 |
| 12 | 3.03E-04 | 3.51E-04 | 1.00E-02 | 7.13E-03 | 3.88E-03 | 3.41E-03 | 3.14E-03 | 9.98E-08 | 1.08E-03 | 1.04E-03 | 3.00E-03 | 3.80E-02 |
| 13 | 4.36E-04 | 6.90E-04 | 1.70E-02 | 1.40E-02 | 4.42E-03 | 8.17E-03 | 7.40E-03 | 1.15E-06 | 3.49E-03 | 3.52E-03 | 1.09E-02 | 7.16E-02 |
| 14 | 4.19E-04 | 5.11E-04 | 1.22E-02 | 1.06E-02 | 4.42E-03 | 5.51E-03 | 5.02E-03 | 6.80E-07 | 2.22E-03 | 2.17E-03 | 7.31E-03 | 5.37E-02 |
| 15 | 4.13E-04 | 4.51E-04 | 1.05E-02 | 9.37E-03 | 4.42E-03 | 4.70E-03 | 4.30E-03 | 4.94E-07 | 1.85E-03 | 1.78E-03 | 6.06E-03 | 4.78E-02 |
| 16 | 4.01E-04 | 3.47E-04 | 7.47E-03 | 7.13E-03 | 4.42E-03 | 3.41E-03 | 3.14E-03 | 1.20E-07 | 1.27E-03 | 1.19E-03 | 3.91E-03 | 3.80E-02 |
| 17 | 5.55E-04 | 6.83E-04 | 1.38E-02 | 1.40E-02 | 4.99E-03 | 8.17E-03 | 7.40E-03 | 1.50E-06 | 4.00E-03 | 3.97E-03 | 1.25E-02 | 7.16E-02 |
| 18 | 5.35E-04 | 5.06E-04 | 9.51E-03 | 1.06E-02 | 4.99E-03 | 5.51E-03 | 5.02E-03 | 8.83E-07 | 2.56E-03 | 2.45E-03 | 8.59E-03 | 5.37E-02 |
| 19 | 5.28E-04 | 4.46E-04 | 8.01E-03 | 9.37E-03 | 4.99E-03 | 4.70E-03 | 4.30E-03 | 6.35E-07 | 2.13E-03 | 2.02E-03 | 7.24E-03 | 4.78E-02 |
| 20 | 5.12E-04 | 3.43E-04 | 5.39E-03 | 7.13E-03 | 4.99E-03 | 3.41E-03 | 3.14E-03 | 1.39E-07 | 1.47E-03 | 1.35E-03 | 4.87E-03 | 3.80E-02 |

*Essential amino acids

**Table S5.** Concentrations of the macronutrients added to the nutribloods

| Nutriblood | Proportion Protein | Conc | Protein (g/L) | Total carbs (g/L) | Lipids (g/L) | Sum of simple sugars (g/L) | Glycogen (g/L) |
|---|---|---|---|---|---|---|---|
| 1 | 0.17 | 63 | 22.841 | 1.66951 | 1.0068 | 0.1768 | 1.49271 |
| 2 | 0.17 | 42 | 19.841 | 1.65672 | 1.0068 | 0.1768 | 1.47991 |
| 3 | 0.17 | 33.6 | 18.750 | 1.65160 | 1.0068 | 0.1768 | 1.47479 |
| 4 | 0.17 | 16.8 | 16.735 | 1.64137 | 1.0068 | 0.1768 | 1.46456 |
| 5 | 0.33 | 63 | 29.468 | 1.42074 | 1.0068 | 0.1768 | 1.24393 |
| 6 | 0.33 | 42 | 25.635 | 1.40794 | 1.0068 | 0.1768 | 1.23114 |
| 7 | 0.33 | 33.6 | 24.240 | 1.40283 | 1.0068 | 0.1768 | 1.22602 |
| 8 | 0.33 | 16.8 | 21.665 | 1.39259 | 1.0068 | 0.1768 | 1.21578 |
| 9 | 0.50 | 63 | 34.976 | 1.15641 | 1.0068 | 0.1768 | 0.97961 |
| 10 | 0.50 | 42 | 30.450 | 1.14362 | 1.0068 | 0.1768 | 0.96681 |
| 11 | 0.50 | 33.6 | 28.803 | 1.13850 | 1.0068 | 0.1768 | 0.96169 |
| 12 | 0.50 | 16.8 | 25.763 | 1.12827 | 1.0068 | 0.1768 | 0.95146 |
| 13 | 0.67 | 63 | 37.541 | 0.89209 | 1.0068 | 0.1768 | 0.71528 |
| 14 | 0.67 | 42 | 32.692 | 0.87930 | 1.0068 | 0.1768 | 0.70249 |
| 15 | 0.67 | 33.6 | 30.927 | 0.87418 | 1.0068 | 0.1768 | 0.69737 |
| 16 | 0.67 | 16.8 | 27.671 | 0.86394 | 1.0068 | 0.1768 | 0.68714 |
| 17 | 0.83 | 63 | 36.625 | 0.64332 | 1.0068 | 0.1768 | 0.46651 |
| 18 | 0.83 | 42 | 31.891 | 0.63052 | 1.0068 | 0.1768 | 0.45371 |
| 19 | 0.83 | 33.6 | 30.168 | 0.62540 | 1.0068 | 0.1768 | 0.44860 |
| 20 | 0.83 | 16.8 | 26.989 | 0.61517 | 1.0068 | 0.1768 | 0.43836 |
